# Identification and Characterization of a B-Raf Kinase Alpha Helix Critical for the Activity of MEK Kinase in MAPK Signaling

**DOI:** 10.1101/2020.07.19.211177

**Authors:** Diep Nguyen, Linda Yingqi Lin, Jeffrey O. Zhou, Emily Kibby, Twan W. Sia, Tiara D. Tillis, Narine Vapuryan, Ming-Ray Xu, Rajiv Potluri, YongJoon Shin, Elizabeth A. Erler, Naomi Bronkema, Daniel J. Boehlmer, Christopher D. Chung, Caroline Burkhard, Shirley H. Zeng, Michael Grasso, Lucila A. Acevedo, Ronen Marmorstein, Daniela Fera

**Affiliations:** Department of Chemistry and Biochemistry, Swarthmore College, Swarthmore, PA 19081; Department of Chemistry, University of Pennsylvania, Philadelphia, PA, 19104; Department of Biochemistry and Biophysics, Perelman School of Medicine, University of Pennsylvania, Philadelphia, PA, 19104; Abramson Family Cancer Research Institute, Perelman School of Medicine, University of Pennsylvania, Philadelphia, PA, 19104

**Author notes:** **Corresponding Author**: Daniela Fera, Swarthmore College, Department of Chemistry and Biochemistry, Swarthmore, PA 19081, United States.

## Abstract

In the MAPK pathway, an oncogenic V600E mutation in B-Raf kinase causes the enzyme to be constitutively active, leading to aberrantly high phosphorylation levels of its downstream effectors, MEK and ERK kinases. The V600E mutation in B-Raf accounts for more than half of all melanomas and ∼3% of all cancers and many drugs target the ATP-binding site of the enzyme for its inhibition. Since B-Raf can develop resistance against these drugs and such drugs can induce paradoxical activation, drugs that target allosteric sites are needed. To identify other potential drug targets, we generated and kinetically characterized an active form of B-Raf^V600E^ expressed using a bacterial expression system. In doing so, we identified an alpha helix on B-Raf, found at the B-Raf-MEK interface, that is critical for their interaction and the oncogenic activity of B-Raf^V600E^. We performed binding experiments between B-Raf mutants and MEK using pull downs and biolayer interferometry, and assessed phosphorylation levels of MEK *in vitro* and in cells as well as its downstream target ERK to show that mutating certain residues on this alpha helix is detrimental to binding and downstream activity. Our results suggest that this B-Raf alpha helix binding site on MEK could be a site to target for drug development to treat B-Raf^V600E^-induced melanomas.

## Introduction

The mitogen-activated protein kinase (MAPK) pathway plays an important role in cellular growth and differentiation through the downstream effects of its activation. As such, its dysregulation leads to a number of cancers, in many cases because protein mutations lead to constitutive activity and uncontrolled cell proliferation ^*1–3*^. This pathway is initiated by growth factor binding to a receptor at the cell membrane, thereby activating a number of adaptor molecules which in turn activate Ras. Active Ras binds to guanosine triphosphate (GTP), which allows it to associate with the Ras binding domain of Raf, a serine/threonine kinase, at the membrane ^*4–6*^. The binding of Ras to the N-terminal domain of Raf causes a conformational change in Raf that frees the C-terminal kinase domain from the N-terminal domain ^*7*^. Raf exists in the cytosol as a quiescent Raf-MEK complex ^*8*^, and Ras binding promotes formation of heterotetramers (MEK—B-Raf—B/C-Raf—MEK). This activation leads to the phosphorylation of MEK1/2, downstream activation of ERK1/2, and cell proliferation at later stages of the MAPK pathway ^*9*^.

Raf kinases have three distinct isoforms in humans – A-Raf, B-Raf, and C-Raf. The B-Raf isoform has been shown to be particularly susceptible to mutation, with over half of melanomas containing activating B-Raf mutations, of which over 90% involve a particular mutant, B-Raf^V600E *10, 11*^. This mutation renders B-Raf constitutively active, independent of both Ras activation and formation of tetrameric complexes. B-Raf^V600E^ mutants are locked in an active conformation because the negatively charged glutamate residue mimics phosphorylation of the activation loop, which releases its inhibitory interaction with the ATP-binding domain, found between the N- and C-lobes of the kinase domain ^*12*^. The B-Raf isoform also has the strongest activity towards MEK ^*13*^. B-Raf^V600E^ mutants mediate unregulated phosphorylation of two conserved residues in the MEK activation loop, S218 and S222 of MEK1 (or S222 and S226 of MEK2) ^*14–16*^. This phosphorylation increases MEK activity, which leads to the binding, phosphorylation, and activation of ERK. ERK1 gets phosphorylated on T202/Y204 and ERK2 gets phosphorylated on T183/Y185 ^*17, 18*^. Activated ERK can translocate to the nucleus to phosphorylate and regulate transcription factors resulting in changes in gene expression ^*19*^. Some ERK substrates are also found in the cytoplasm and other organelles ^*20*^.

Drug discovery efforts against oncogenic B-Raf, mainly B-Raf^V600E^, have largely focused on targeting the ATP-binding pocket and several (e.g. vemurafenib and dabrafenib) have achieved approval by the FDA for treatment of some B-Raf-mutated melanomas ^*10, 21, 22*^. However, acquired resistance to these competitive inhibitors and paradoxical activation in cells harboring wild-type B-Raf or mutants other than B-Raf^V600E^ show the need for allosteric inhibitors and combination therapies ^*23–26*^. The B-Raf-MEK interface represents a potential target for allosteric inhibition of B-Raf activity as this binding interaction is critical for the phosphorylation of MEK and the activation of downstream targets ^*23, 27*^. Mutational studies using the available crystal structure of the complex can provide site-specific information about particular surfaces that may be promising drug targets due to their role in enhancing or inhibiting the B-Raf-MEK interaction ^*8*^.

Many studies of B-Raf rely on the use of Sf9 insect cells for expression since it has been well established to produce the active enzyme ^*28*^. Mammalian cells have also been used, but yields are often low ^*29, 30*^. A recent report had modified the B-Raf kinase domain by the introduction of 16 mutations to improve soluble protein expression using bacterial cells for X-ray crystallographic studies ^*31*^. Unfortunately, expression of the B-Raf^V600E^ kinase domain with these 16 surface mutations (called B-Raf^V600E,16mut^) yielded an inactive enzyme despite no mutations of key catalytic residues, and the ability to bind ATP was intact (though its K_M_ was not determined). Using B-Raf^V600E,16mut^ expressed in bacteria as a starting point, we generated an active B-Raf construct that can be produced in *E. coli*. We further identified and characterized an α-helix (residues 661-671) in B-Raf that we showed is critical for the interaction with MEK. This site of interaction offers a new target for therapeutics against B-Raf^V600E^-induced malignancies.

## Materials and Methods

### Expression and Purification of Proteins

DNA encoding the B-Raf kinase domain residues 448-723 containing 16 solubilizing mutations in the C-lobe (I543A, I544S, I551K, Q562R, L588N, K630S, F667E, Y673S, A688R, L706S, Q709R, S713E, L716E, S720E, P722S, and K723G) for protein expression in *E. coli* was ordered from Epoch Biolabs. This construct was cloned into a pRSF vector with an N-terminal GST tag and a TEV protease cleavage site between the tag and protein. Site-directed mutagenesis was performed using the manufacturer’s protocols (Stratagene) to introduce mutations. GST-tagged B-Raf^V600E,16mut^ and B-Raf^V600E,15mut^ proteins were expressed in BL21(DE3)RIL cells at 37 °C, induced with 1mM IPTG overnight at 16 °C, spun down the next day, and lysed in lysis buffer (50 mM potassium phosphate pH 7.0, 250 mM NaCl, DNase I, 1 mM PMSF). The lysate was spun down at 19,000 rpm for 30 minutes and the supernatant was incubated with GST-Bind resin (Millipore) at 4 °C for 1 hr. The protein on the resin was washed with wash buffer (50 mM potassium phosphate pH 7.0, 250 mM NaCl) overnight and eluted the next morning in wash buffer with 20 mM glutathione and 5 mM βME. The protein was concentrated using a 30 kDa cutoff centrifugal filter unit (Millipore) and chromatographed on a Superdex 200 increase column in a final buffer of 20 mM HEPES pH 7.0, 150 mM NaCl, 5% glycerol and 5 mM βME. Protein eluted as a mixture of dimer and monomer and both species were pooled together, concentrated and flash frozen in liquid nitrogen. For certain experiments, the GST-tag was cleaved off. To do this, the protein was not eluted off the GST-Bind resin. Instead, TEV protease was added to the column and incubated overnight at 4 °C. The next morning, the cleaved protein was washed off with 50 mM potassium phosphate pH 7.0, 25 mM NaCl and run over both SP Sepharose and Q Sepharose ion exchange columns in tandem and the flow through was collected. The protein was then concentrated using a 10 kDa cutoff centrifugal filter unit (Millipore) and chromatographed on a Superdex 200 gel filtration column in a final buffer of 20 mM HEPES pH 7.0, 150 mM NaCl, 10 mM DTT and 5% glycerol. Protein eluted as a mix of dimer and monomer and both species were pooled together, concentrated and flash frozen in liquid nitrogen. Freeze-thawing reduced enzyme activity, so each aliquot was thawed only once.

DNA encoding the B-Raf kinase domain residues 442-724 was used as a template and cloned into a Pfastbac dual vector for protein expression in baculovirus infected Sf9 insect cells. An N-terminal 6x-His tag was inserted into the plasmid, and this plasmid was used as a template to create the mutant B-Raf^V600E^ using site-directed mutagenesis according to the manufacturer’s protocols (Stratagene). His-tagged B-Raf^V600E^ kinase domain was expressed in insect cells essentially as previously described ^*22*^. Briefly, protein constructs were coexpressed with full length mouse p50^cdc37^ as an expression chaperone, pelleted, suspended in lysis buffer 2 (25 mM pH Tris 8.0, 250 mM NaCl, 5 mM imidazole, and 10% glycerol) treated with Complete EDTA-free protease inhibitor cocktail tablets (Roche) and DNase I. After lysis, cell lysates were centrifuged at 19,000 rpm and then incubated with TALON metal affinity resin for 1 hr at 4 °C. The resin was then washed extensively with lysis buffer 2 and the protein was eluted with 25 mM Tris pH 7.5, 250 mM NaCl, 250 mM imidazole, and 10% glycerol. The protein was then diluted in low salt buffer containing 1 mM EDTA and 1 mM DTT before being run on an SP Sepharose column. The protein was eluted with a salt gradient from 0 mM NaCl to 1 M NaCl. Peak fractions were pooled and run on a Superdex 200 gel filtration column and stored in a final buffer of 25 mM Tris pH 8.0, 300 mM NaCl, 1 mM DTT and 10% glycerol. Protein was concentrated to ∼0.5 mg/mL and flash frozen in liquid nitrogen. Freeze-thawing reduced enzyme activity, so each aliquot was thawed only once.

A cDNA library for full length MEK was purchased from Dharmacon (Catalog # MHS 6278-211690391). Full length human MEK1 was cloned into a Pet28a(+) vector encoding an N-terminal 6x-His tag and a TEV protease cleavage site between the tag and protein. MEK with an N-terminal 6x-His tag was expressed in DE(3) RIL bacterial expression cells at 37 °C, induced with 1 mM IPTG overnight at 16 °C, spun down the next day, and lysed in lysis buffer 3 (25 mM Tris pH 7.5, 500 mM NaCl, 5 mM βME) supplemented with 1 mM PMSF and DNase I. The lysate was spun down at 19,000 rpm for 30 minutes and the supernatant was added to Ni-NTA metal affinity resin (Thermo Scientific) and left to incubate at 4 °C for 1 hr. The supernatant was then eluted via gravity column and the resin was washed with lysis buffer 3 supplemented with 20 mM imidazole. The MEK protein was eluted with lysis buffer 3 supplemented with 300 mM imidazole. The eluent was dialyzed into dialysis buffer (25 mM Tris pH 7.5, 25 mM NaCl, 5 mM βME) overnight and then applied to a 5 mL Q Sepharose cation exchange column followed by washing in dialysis buffer and elution in 25 mM Tris pH 7.5, 5 mM βME, and 1 M NaCl. Peak fractions were run on an SDS-PAGE gel, pooled, concentrated using a 30 kDa cutoff centrifugal filter unit (Millipore), and applied to a Superdex 200 increase column in a buffer of 20 mM HEPES pH 7.0, 150 mM NaCl, 5% glycerol and 5 mM βME. Protein was concentrated, flash frozen in liquid nitrogen, and stored at −80 °C. Freeze-thawing reduced enzyme activity, so each aliquot was thawed only once.

Full length human MEK1 with an N-terminal GST fusion tag and a C-terminal His tag in a pGex-3t vector was provided by Dr. Michael Olson (Beatson Institute for Cancer Research, Glasgow, UK). GST-MEK1 fusion protein used as substrate in ELISA assays was prepared essentially as previously described ^*22*^. Briefly, protein was expressed in BL21(DE3)RIL cells at 37 °C and induced with 0.5 mM IPTG at 15 °C overnight. The cells were then harvested and resuspended in lysis buffer 4 (20 mM HEPES pH 7.5, 500 mM NaCl, 10 mM βME, 10 mM imidazole, 5% glycerol) supplemented with 1 mM PMSF and DNase I. The lysate was sonicated and spun down, and the supernatant was added to Ni-NTA resin and incubated for 1 hr at 4 °C. The resin was then washed extensively with lysis buffer 4 and eluted with lysis buffer 4 supplemented with 250 mM imidazole. Peak fractions were then concentrated using a 50k Dalton cutoff centrifugal filter unit (Millipore) and loaded onto a Superdex 200 gel filtration column in a final buffer of 20 mM HEPES pH 7.0, 150 mM NaCl, 10 mM βME, and 5% glycerol. The protein eluted off the sizing column in two separate populations, and the second peak corresponding to a GST-MEK dimer was collected, concentrated to ∼14 mg/mL, flash frozen in liquid nitrogen and stored at −80 °C. Freeze-thawing reduced enzyme activity, so each aliquot was thawed only once.

pGEX6P1-GST-HA-ERK2^K54^ was provided by Donita Brady (University of Pennsylvania). GST-ERK2^K54R^ was expressed in DE(3) RIL cells in TB (Dot Scientific) at 37 °C until reaching an OD_600_ of 0.700 and then induced with 1 mM IPTG overnight at 17 °C. The cells were then spun down the next day and lysed in lysis buffer 5 (25 mM Tris pH 7.5, 500 mM NaCl, and 5 mM βME) treated with 1 mM PMSF and DNase I. The lysate was then spun down at 19,000 rpm for 40 minutes and the supernatant was added to glutathione agarose resin (Pierce product # 16102BID) previously washed with lysis buffer 5 and left to incubate at 4 °C for 2 hr. The supernatant was eluted via gravity column and the resin was then washed with 1 L of lysis buffer 5. The protein was then eluted with lysis buffer 5 supplemented with 20 mM glutathione (Sigma Aldrich, product # G4521) and the elution fractions were pooled together and dialyzed into dialysis buffer (25 mM Tris pH 7.5, 150 mM NaCl, 5 mM βME). The protein was then concentrated and applied to a Superdex 200 10/300 gel filtration column in a final buffer of 25 mM Tris pH 7.5, 150 mM NaCl, 5 mM βME. Protein was concentrated, flash frozen in liquid nitrogen, and stored at −80 °C for future use. Freeze-thawing reduced enzyme activity, so each aliquot was thawed only once.

### Biolayer Interferometry

Kinetic measurements of B-Raf binding to full length MEK were carried out using the fortéBIO BLItz instrument. After an initial baseline in 20 mM HEPES pH 7.0, 150 mM NaCl, 5% glycerol and 5 mM βME, His-tagged MEK or His-tagged B-Raf at a concentration of 0.2 mg/mL was immobilized on Dip and Read™Ni-NTA Biosensors until saturation was reached (2.5 min). GST-B-Raf constructs or GST-MEK were tested at concentrations ranging from 0.25 µM to 20 µM, depending on the particular construct used and its solubility, as well as based on its binding affinity to MEK or B-Raf, respectively. Association and dissociation were each monitored for 3 minutes. Step corrections, at the start of association and at the start of dissociation, were performed manually using Microsoft Excel. Analyses were then performed using a global fit of at least three measurements using nonlinear regression curve fitting to a A + B <=> C model in GraphPad Prism, version 8 to obtain k_on_, k_off_ and K_D_ values.

### Pull Downs

20 µg of GST-MEK was incubated with 40 µL of GST-bind resin in a buffer of 20 mM HEPES pH 7.0, 150 mM NaCl, 5 mM βME, and 0.05% Tween 20 for 15 minutes at 4 °C with gentle agitation. Then, an equimolar amount of the respective B-Raf construct was added and incubated at 4 °C for one hour with gentle agitation. The beads were then washed three times with 1 mL buffer and subjected to SDS-PAGE analysis.

### Western Blotting Based Kinase Assay

Reactions were set up with 1 mM ATP and kinase buffer (100 mM Tris, pH 7.5, 10 mM MgCl_2_ and 5 mM βME) and the indicated kinases. Final concentrations of 50 nM, 200 nM and 200 nM were used for B-Raf, MEK and ERK, respectively. Kinases were diluted in 20 mM HEPES pH 7.0, 150 mM NaCl, 5% glycerol and 5 mM βME prior to addition to reaction mixtures. Phosphorylated MEK and ERK were produced by incubating the enzymes with ATP for 15 min at room temperature. In preliminary experiments, it was determined that 15 minutes was more than sufficient to yield complete activation.

Reactions were quenched with 4x loading dye, subjected to SDS-PAGE and then transferred to nitrocellulose membrane to be visualized by western blotting. Membranes were blocked with 5% (w/v) nonfat milk overnight at 4 °C with gentle rocking. The membranes were then incubated overnight at 4 °C with either rabbit anti-p-MEK antibody against Ser217/Ser221 (1:1000) (Cell Signaling, #9121), rabbit anti-p-ERK antibody against Thr202/Tyr204 (1:1000) (Cell Signaling, #9101), anti-MEK antibody (1:1000) (Cell Signaling, #9122), anti-ERK antibody (1:1000) (Cell Signaling, #9102), or anti-GST antibody (1:1000) (Cell Signaling, #2625). The next day, the membranes were incubated with goat anti-rabbit antibody conjugated to HRP (1:500 or 1:1000 against MEK or ERK/GST, respectively) (BioRad) for one hour at room temperature. Bands were visualized by incubating with a solution containing the HRP substrate, 4CN, a color developing solution, following the manufacturer’s protocol. Quantitative analyses of the immunoblots were executed with ImageJ software.

### Cell-Based Assay

The pCDNA™4/TO and pcDNA3.1-Hygro plasmids containing full length wild-type B-Raf and MEK, respectively, were generously provided by Dr. Zhihong Wang (University of Sciences). Site-directed mutagenesis was performed using the manufacturer’s protocols (Stratagene) to introduce mutations into B-Raf to generate the V600E (B-RAF^FL,V600E^) and V600E, I666R (B-RAF^FL,V600E,I666R^) single and double mutants, respectively. To assess phosphorylation levels in HEK 293T cells, B-Raf and MEK DNA were transiently transfected (in a 1:1 ratio) using linear polyethylenimine (PEI) following the manufacturer’s suggested protocol. After 24 hr of expression, the cells were dislodged from the flasks by gentle tapping, counted and normalized across all samples. The supernatant was then clarified by centrifugation at 3000 RPM at 4 °C. Cells were washed with cold PBS and centrifuged again at 4 °C. The cells were then lysed in RIPA buffer supplemented with a Complete EDTA-free protease inhibitor cocktail tablet (Roche) and Halt protease. Cell debris was pelleted by another round of centrifugation at 4 °C. The supernatant was then subject to Western analysis as described above, using rabbit anti-p-MEK antibody against Ser217/Ser221 (1:500) (Cell Signaling, #9121), rabbit anti-p-ERK antibody against Thr202/Tyr204 (1:1000) (Cell Signaling, #9101), anti-MEK antibody (1:250) (Cell Signaling, #9122), anti-ERK antibody (1:1000) (Cell Signaling, #9102), or anti-B-Raf antibody (1:2000) (Cell Signaling, #D9T6S). Goat anti-rabbit antibody conjugated to HRP (BioRad) was used at 1:500 or 1:1000 against MEK or B-Raf/ERK, respectively.

### ELISA Assay

Kinetics of kinase activity to determine steady state parameters for the ATP substrate of different constructs (Sf9 expressed His-B-Raf^V600E^ mutant and *E. coli* expressed B-Raf^V600E,15mut^) were determined using an assay modified from that previously described ^*22*^. Briefly, GST-MEK fusion protein was diluted 3:1000 (to 600 nM) in Tris-Buffered Saline treated with 0.05% Tween 20 (TBST) and added to a glutathione-coated 96 well plate (Pierce #15240) and incubated at room temperature for 1 hour with shaking. Glutathione-coated plates were washed extensively, and 50 µL of B-Raf (at 50 nM or serial dilutions were performed to evaluate enzyme concentration dependence) in 50 mM HEPES pH 7.0 and 50 mM NaCl was then added to the plate with 50 µL of ATP solution (200 µM ATP or varied ATP concentration to assess K_M_(ATP) and k_cat_) in a buffer containing 50 mM HEPES pH 7.0, 200 mM NaCl, and 20 mM MgCl_2_. The plate was then incubated at RT for 30 minutes to assess concentration of enzyme dependence, for 15 minutes to assess K_M_(ATP) and k_cat_, or quenched with 100 mM EDTA at various time points to determine dependence on reaction time. Two rows of the plate were used to generate a standard curve to determine the concentration of the p-MEK product. To do this, various MEK concentrations were added to the wells. After washing, 50 µL of 50 nM B-Raf was added to the plate with 50 µL of 200 µM ATP solution and incubated at RT for 30 minutes. After incubation with B-Raf and ATP, the reaction was then discarded, and the plate was extensively washed with TBST + 100 mM EDTA and a 1:8,000 dilution of primary antibody (p-MEK1/2 (S217/S221) Rabbit Antibody (Cell Signaling)) was added to the plate and incubated for 1 hour. The plate was then washed with TBST and then incubated with a 1:10,000 dilution of secondary antibody (Goat Anti-Rabbit IgG (H+L)-HRP (BioRad)) for 1 hour. The plate was then washed three times and Supersignal ELISA Pico Chemiluminescent Substrate (Pierce #37069) was added and the plate was read on Promega GloMAX 96 Microplate Luminometer. Each curve was repeated in triplicate, using the standard curve to determine the concentration of reaction product and the time of reaction to determine [Product]/min. GraphPad Prism 5.0 was used to plot and fit with the Enzyme kinetics – Michaelis-Menten model in the Analyze menu.

Kinetics of kinase activity to determine steady state parameters for MEK substrate of different constructs (Sf9 expressed His-B-Raf^V600E^ mutant and *E. coli* expressed B-Raf^V600E^ mutants) were determined by an ELISA assay. Briefly, GST-HA-ERK2^K54^ protein was diluted 1:10 (to 3.26 uM) in Tris-Buffered Saline treated with 0.05% Tween 20 (TBST). The diluted ERK was added to a glutathione-coated 96 well plate (Pierce #15240) and incubated at room temperature for 1 hour with shaking. The various GST-tagged B-Raf mutants were diluted in 50 mM HEPES pH 7.0 and 50 mM NaCl so that the final concentration of B-Raf in the combined reaction was 25 nM. An equal volume of His-MEK was incubated with each B-Raf mutant at 0 µM − 2.2 µM final concentrations in the combined reaction for 1 hour with shaking in a separate plate. The glutathione-coated plates were then washed extensively, and 50 µL of B-Raf-MEK complex solution was then added to the plate with 50 µL of 200 µM ATP in a buffer of 50 mM HEPES pH 7.0, 200 mM NaCl, and 20 mM MgCl_2_. Two rows of the plate were used to generate a standard curve to determine the concentration of the p-ERK product. To do this, various concentrations of GST-HA-ERK2^K54^ were added to the wells. After washing, 50 µL of B-Raf-MEK complex (formed by mixing GST-B-Raf and MEK at a final concentration of 133.6 nM and 289.3 uM, respectively) was added to the plate with 50 µL of 200 µM ATP. The plate was incubated at RT for 15 minutes, the reaction was discarded, and the plate was extensively washed with TBST + 100 mM EDTA. A 1:5,000 dilution of primary antibody (p-ERK (Thr202/Tyr204) Rabbit Antibody (Cell Signaling)) was added to the plate and incubated for 1 hour. The plate was then washed two times with TBST and then incubated with a 1:5,000 dilution of secondary antibody (Goat Anti-Rabbit IgG (H+L)-HRP (BioRad)) for 1 hour. The plate was then washed three times and Supersignal ELISA Pico Chemiluminescent Substrate (Pierce #37069) was added and the plate was read on a Promega GloMAX 96 Microplate Luminometer. Each curve was repeated in triplicate, and GraphPad Prism 5.0 was used to plot and fit with the Enzyme kinetics – Michaelis-Menten model in the Analyze menu.

## Results

### Restoration of MEK Complex Formation with the Solubilized *E. coli* Construct of B-Raf

The 16 solubilizing mutations introduced into B-Raf (B-Raf^V600E,16mut^) for bacterial expression are located on the surface of the C-lobe of the kinase domain (Figure 1A, 1B). Interestingly, the combination of these mutations rendered B-Raf inactive, even with the V600E mutation, as it was unable to bind to and phosphorylate MEK. Analysis of the B-Raf-MEK complex structure ^*8*^ revealed that the Phe at position 667 could make a key hydrophobic interaction with MEK, and therefore the solubilizing mutation to Glu could have led to charge-charge repulsion with the acidic patch of MEK in which D315 resides (Figure 1C). To test this hypothesis, we generated a mutant in which E667 was reverted back to Phe (B-Raf^V600E,15mut^) for binding and activity assays.

**Figure 1.**
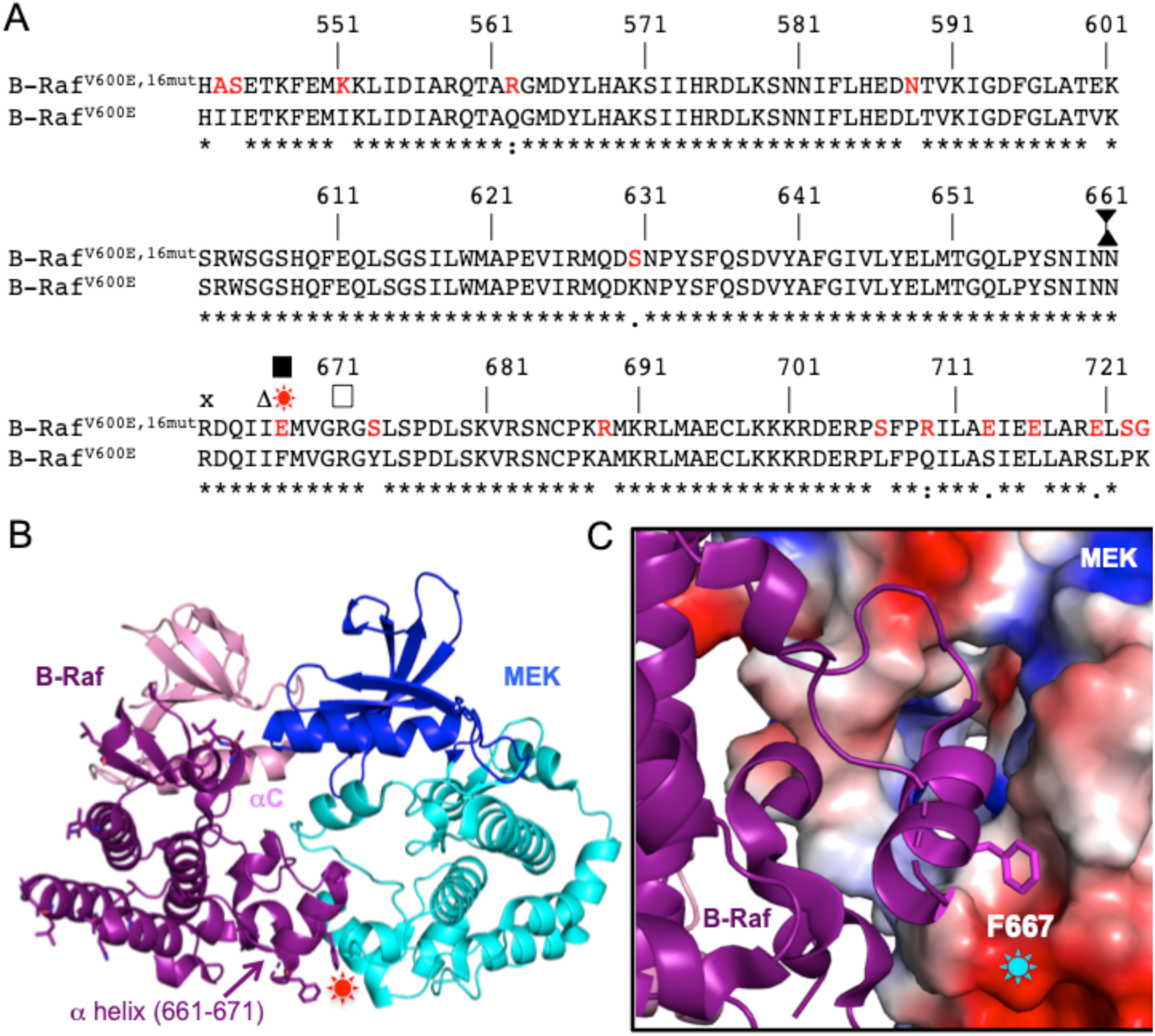
B-Raf-MEK Complex. (A) Sequence alignments of B-Raf kinase domain C-lobe. Mutations in the solubilized construct are shown in red. Mutations introduced in this study are shown with various symbols. Conserved residues with respect to wild-type B-Raf with the V600E mutation are marked by an asterisk below the alignment. (B) Structure of the B-Raf-MEK complex. The two proteins are shown as cartoons with the B-Raf N-lobe in pink, B-Raf C-lobe in purple, MEK N-lobe in blue, and MEK C-lobe in cyan. The residue that was mutated to produce active B-Raf with 15 surface mutations is shown in sticks and highlighted with a red star. The α-helix under study is indicated and the αC helix is labeled. Figure generated using PDB ID: 4MNE ^*8*^. (C) Zoomed in view of the B-Raf α-helix. The α-helix is shown as a purple cartoon with the solubilizing mutation that interacts with MEK shown as sticks. The electrostatic surface potential of the MEK C-lobe is shown (blue is positively charged and red is negatively charged) and the position of D315 is shown with a cyan star. Figure generated using PDB ID: 4MNE ^*8*^.

This mutant was expressed in *E. coli* with either a 6x-His or GST tag. The size-exclusion chromatography (SEC) elution profile of the GST-tagged B-Raf^V600E,15mut^ (MW ∼60 kDa) showed two peaks (Figure 2). SDS-PAGE analysis demonstrated that both peaks contain B-Raf; however, absorbance readings demonstrate a high 260/280nm ratio for the first peak. Together with a comparison to standards, this suggests that the first peak contained aggregates. Moreover, the second peak has minimal contamination of other proteins and has dimeric B-Raf (the V600E activating mutation induces B-Raf dimerization in the absence of the N-terminal regulatory domain^*32*^), so fractions from this peak were kept for further analysis.

**Figure 2.**
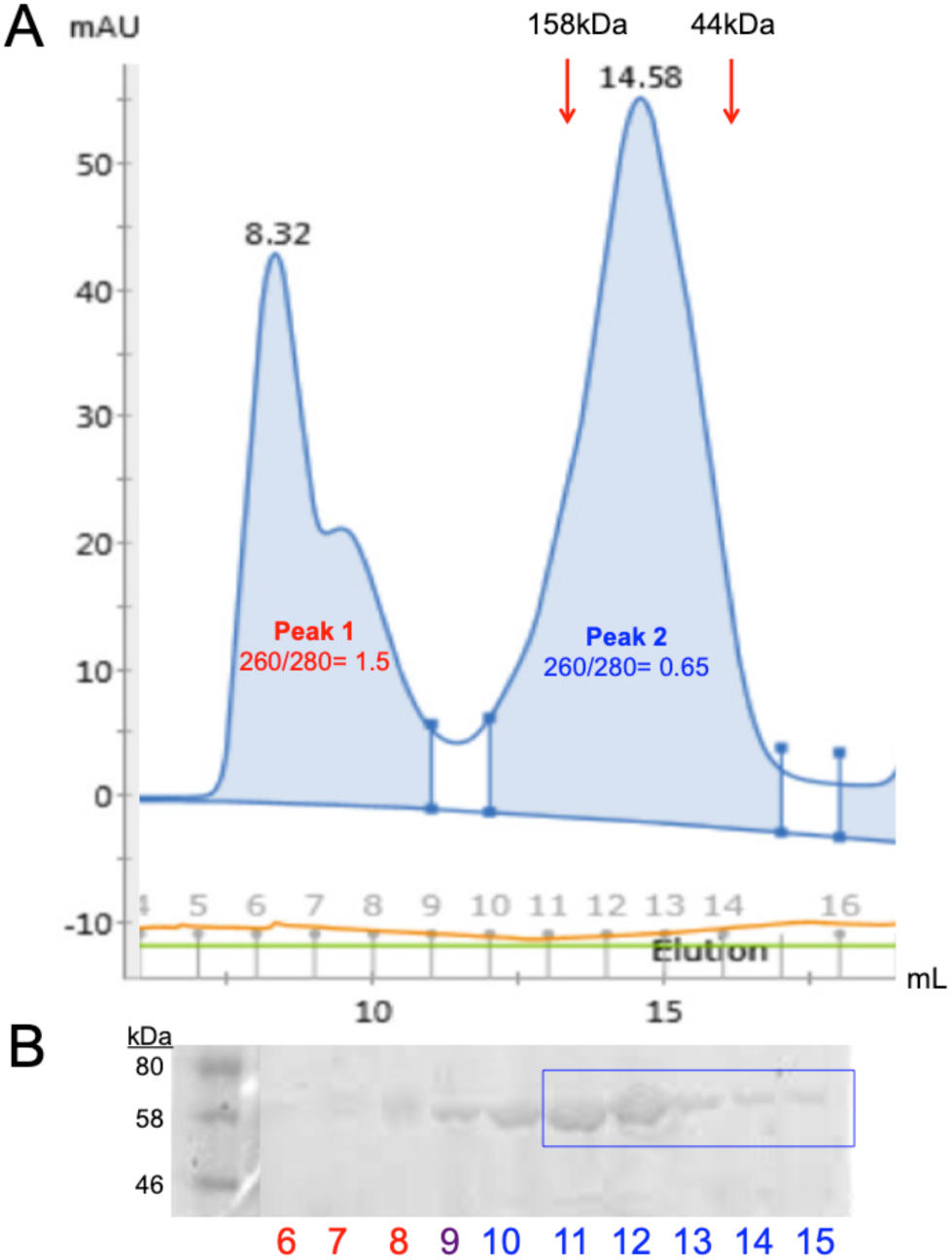
Purification of GST-B-Raf^V600E,15mut^ from *E. coli*. (A) FPLC trace of size-exclusion chromatography run of GST-B-Raf ^V600E,15mut^. The elution volumes of protein standards (158 and 44 kDa) are indicated by arrows. (B) SDS-PAGE analysis of fractions from the corresponding size-exclusion chromatography run. Fraction numbers are indicated and those kept for analysis are boxed.

To assess the ability of B-Raf^V600E,15mut^ to bind MEK and compare it to B-Raf^V600E^, we performed pull-downs. GST-tagged full length wild-type MEK was immobilized on beads and binding of the kinase domain of different B-Raf mutants was visualized using SDS-PAGE. Leaving E667 in the *E. coli* construct ablates complex formation between B-Raf ^V600E,16mut^ and MEK, whereas the E667F reversion restored B-Raf-MEK complex formation *in vitro*, to a similar extent as with B-Raf^V600E^ (Figure 3A). To quantify the binding affinity of B-Raf to MEK, we used biolayer interferometry (BLI). Either His-tagged MEK was immobilized and binding of GST-B-Raf was tested (Figure 3B, left panels), or His-tagged B-Raf was immobilized and binding of GST-MEK was tested (Figure 3B, right panels). Consistent with the results from pull-downs, no binding of GST-B-Raf^V600E,16mut^ was detected, but GST-B-Raf^V600E,15mut^ binding to MEK could be detected with a K_D_ of ∼32 nM (Table 1). This binding affinity is ∼15-fold higher than that of the wild-type B-Raf produced from Sf9 insect cells (K_D_∼481 nM), but similar to that obtained for B-Raf^V600E^ (K_D_ of ∼56 nM). These data suggest that the E667F reversion restored the binding ability of the *E. coli* construct B-Raf^V600E,15mut^ to MEK to what is observed for B-Raf^V600E^ produced in Sf9 insect cells.

**Table 1.**
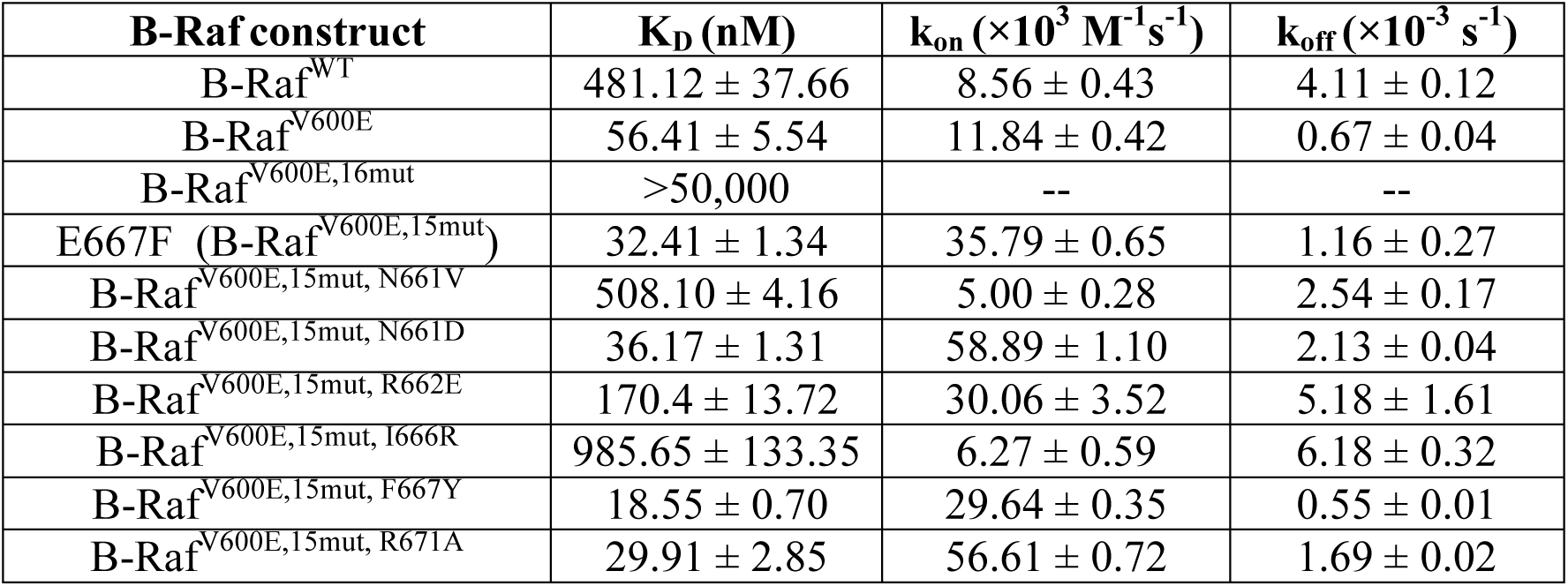
Equilibrium dissociation constant (K_D_) and association and dissociation rate constants (k_on_ and k_off_) from BLI for indicated B-Raf kinase domain constructs and full length wild-type MEK

**Figure 3.**
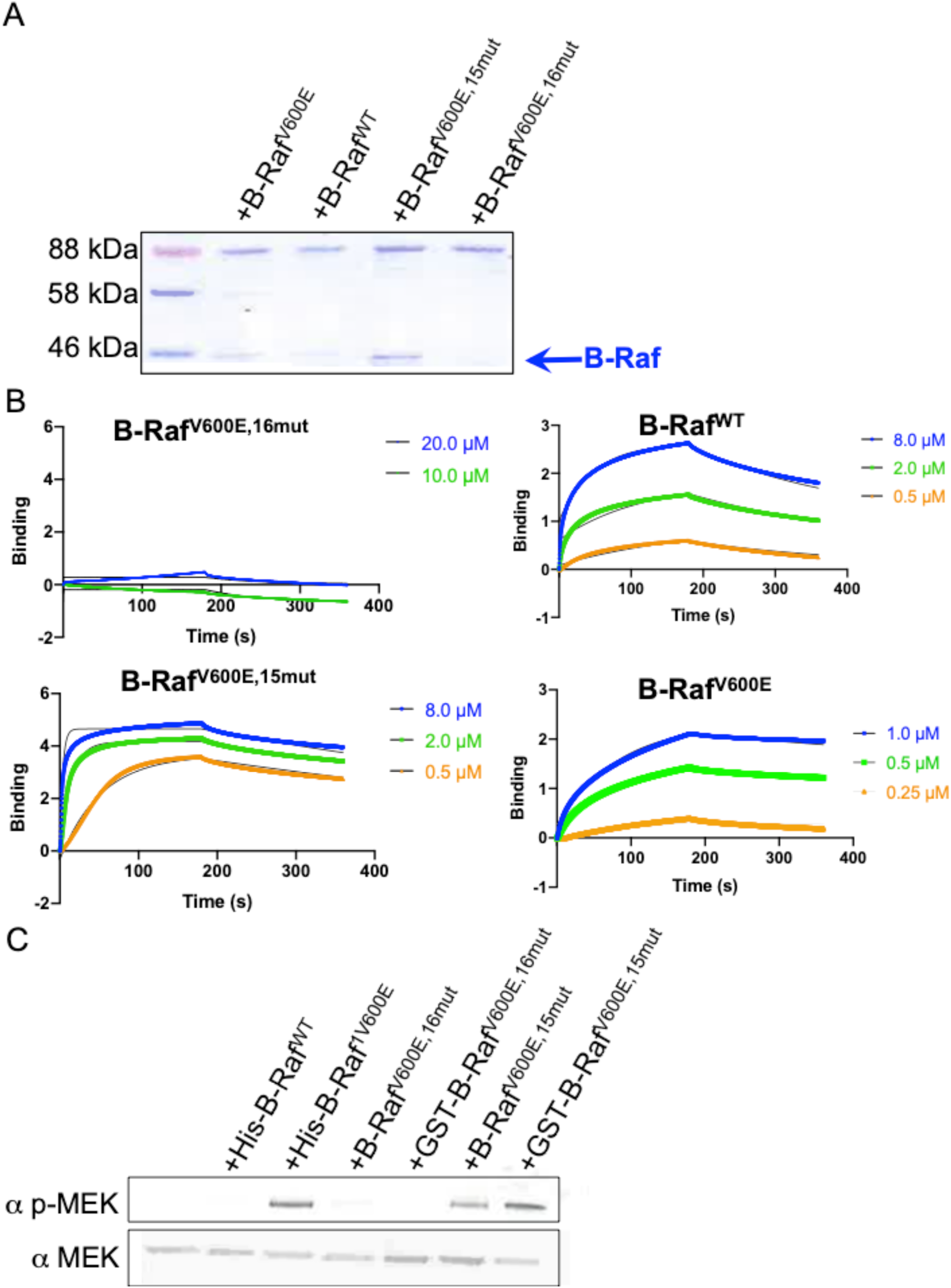
Binding and activity of *E. coli* expressed B-Raf^V600E,15mut^ compared to Sf9-expressed B-Raf^V600E^. (A) SDS-PAGE gel of pull-down assay of GST-tagged full length wild-type MEK with B-Raf^V600E,16mut^ or B-Raf^V600E,15mut^ expressed in *E. coli*, B-Raf^WT^ expressed in Sf9 cells and B-Raf^V600E^ expressed in Sf9 cells. The B-Raf band is indicated with a blue arrow. (B) BLI curves of GST-B-Raf ^V600E,16mut^, GST-B-Raf ^V600E,15mut^, His-B-Raf^WT^, and His-B-Raf^V600E^ binding to full length MEK. In each case of GST-tagged B-Raf mutants, His-tagged MEK was immobilized onto a Ni-NTA biosensor, and B-Raf constructs were introduced at two or more concentrations, ranging from low micromolar to high micromolar, depending on the mutant tested. In each case of His-tagged B-Raf mutants, B-Raf was immobilized onto a Ni-NTA biosensor, and GST-MEK was introduced at least three concentrations, ranging from low micromolar to high micromolar, depending on the mutant tested. Global fits are shown as black lines. (C) A representative Western blot for *in vitro* phosphorylation of MEK, performed in duplicate, using His-tagged B-Raf^V600E^ and B-Raf^WT^ expressed in Sf9 cells compared to B-Raf^V600E,16mut^ and B-Raf ^V600E,15mut^ expressed in *E. coli* with 1 mM ATP.

### MEK can be Phosphorylated by B-Raf^V600E,15mut^

Given that B-Raf^V600E,15mut^ could bind to MEK, and because different expression systems could generate proteins with distinct post-translational modifications and therefore distinct activity levels, we next wanted to determine if B-Raf^V600E,15mut^ could phosphorylate MEK, its downstream target, and if it does so to the same extent and reaction kinetics as B-Raf^V600E^ produced in Sf9 insect cells. To test this, we incubated B-Raf^V600E,15mut^ (either untagged or with a GST tag) or 6x-His-tagged B-Raf^V600E^ together with MEK and ATP and probed for phosphorylation at MEK-Ser218/222 after 15 minutes (Figure 3C). This revealed a similar level of phosphorylated MEK by the V600E containing B-Rafs (B-Raf^V600E,15mut^ produced in *E. coli*, and B-Raf^V600E^ produced in Sf9 insect cells), and a higher level than what was observed by the wild-type B-Raf produced in Sf9 insect cells.

We then used an ELISA assay for a quantitative analysis of the kinase activities of these two V600E-containing B-Raf constructs. We first determined steady state conditions where the reaction rate is linear versus reaction time and enzyme concentration for each B-Raf we tested (Figure S1A, S1B, S1D and S1E). We chose to work with B-Raf at a concentration of 25 nM and a reaction time of 15 minutes to follow steady-state kinetics. The K_M_ of B-Raf^V600E,15mut^ for ATP was measured to be 27.81 **±**6.64 µM (Figure S1C) and of B-Raf^V600E^ for ATP was 0.57 **±**0.17 µM (Figure S1F). These studies revealed significant B-Raf^V600E,15mut^ phosphorylation activity, with apparent Michaelis-Menten kinetics. To obtain K_M_(MEK) for B-Raf^V600E^ and B-Raf^V600E,15mut^, we varied the MEK concentration in the presence of 25 nM B-Raf and high concentrations of ERK and ATP. K_M_(MEK) for B-Raf^V600E^ and B-Raf^V600E,15mut^ are both low-nanomolar (Table 2). This similarity is consistent with the data from Western analysis above and with the K_D_ values obtained from BLI. Together, these data suggest that we were able to restore the activity of the mutated *E. coli* produced protein.

**Table 2.**
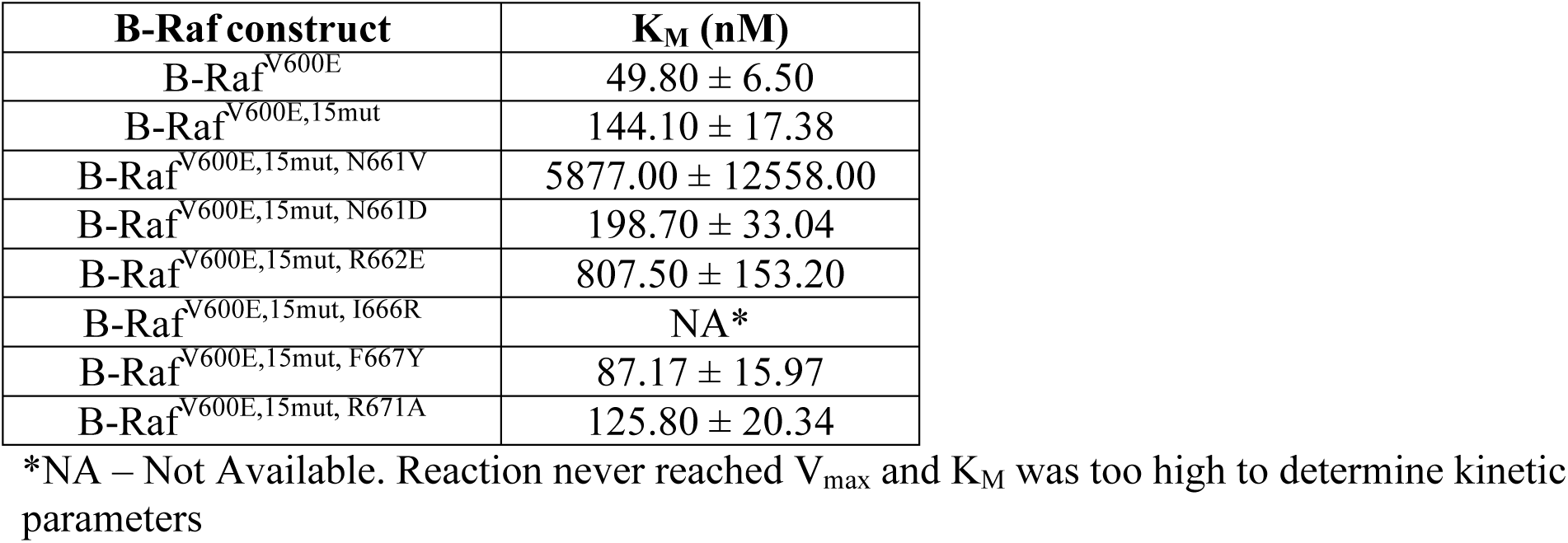
Michaelis Menten constants (K_M_(MEK)) from ELISA for indicated B-Raf kinase domain constructs and full length wild-type MEK

### A B-Raf α-Helix is Critical for Binding MEK and for MEK Phosphorylation

Reversal of the F667E mutation back to F667 was shown to restore the active state of B-Raf, suggesting an important role for this residue and the α-helix on which it resides in B-Raf-MEK complex formation. To verify the importance of this section of the α-helix for complex formation, we further mutated F667 to a Tyr to see if that would enhance the activity of B-Raf. We reasoned that this mutation may introduce a hydrogen bond with the nearby D315 and/or N319 of MEK (Figure 4A) while preserving interactions with the MEK hydrophobic pocket. We tested binding by BLI and found that the introduction of the F667Y mutation in B-Raf^V600E,15mut^ enhanced binding affinity to MEK by almost two-fold, due to a reduced dissociation rate of MEK from B-Raf (Figure 4B,Table 1).

**Figure 4.**
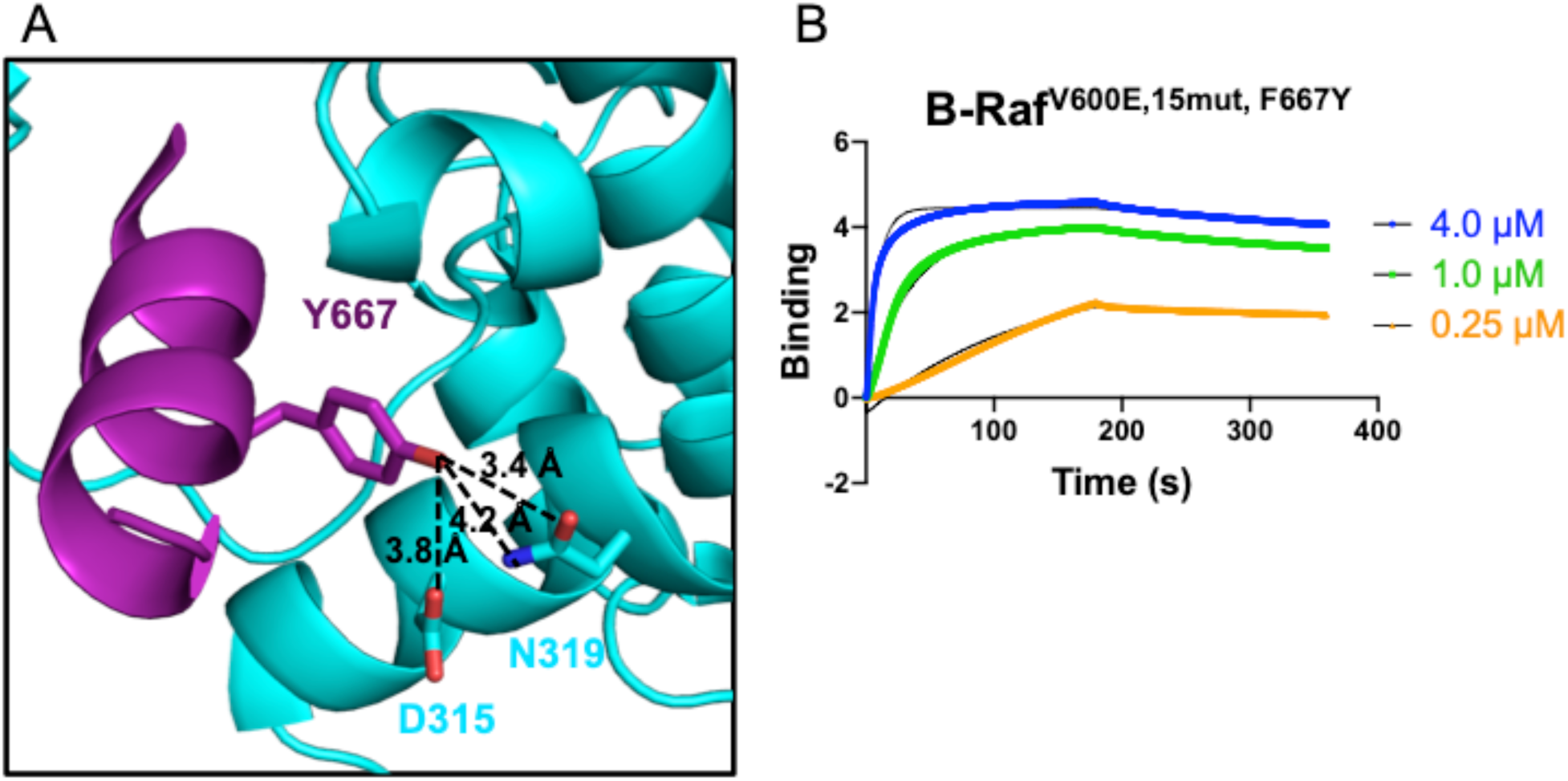
Improving B-Raf binding to MEK. (A) The introduced B-Raf Y667 mutation was modeled using PyMOL by keeping the same coordinates of the backbone and Cβ atoms and is shown as purple sticks together with its possible interactions with MEK residues (cyan). Hydrogen bond distances are shown with dashed lines and distances are provided. Figure generated using PDB ID: 4MNE. ^*8*^ (B) Representative BLI curves of the GST-B-Raf^V600E,15mut,F667Y^ mutant binding to immobilized full length His-tagged MEK. B-Raf was introduced at the indicated concentrations. Global fits are shown as black lines.

The α-helix on which residue 667 resides reaches into a tunnel in MEK kinase (Figure 5A), so we next wanted to determine if the rest of the α-helix (residues 661-671) is critical for complex formation with MEK. To test this, we generated mutations along this α-helix and determined their effects on MEK binding. Specifically, we produced B-Raf^V600E,15mut^ with point mutations N661V, R662E, I666R, and R671A to try to disrupt MEK binding based on the hydrogen bond interactions and complementary electrostatics observed around these residues (Figure 5B-5D). The N661D point mutation was also introduced to try to improve the electrostatic interaction with MEK (Figure 5B).

**Figure 5.**
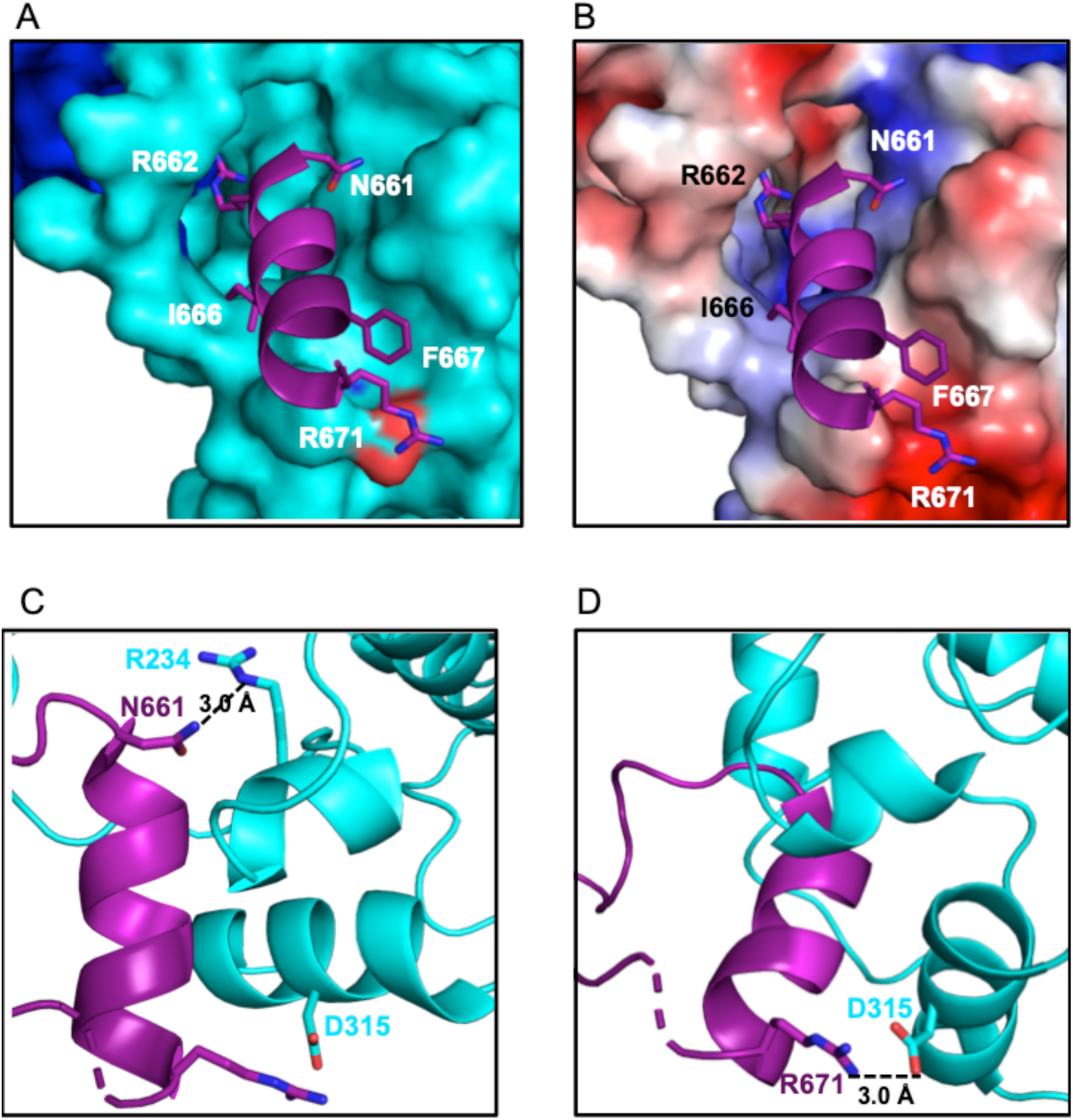
Interactions of B-Raf α-helix with the MEK C-lobe. (A) The B-Raf α-helix under study is shown as a purple cartoon with residues mutated in this study shown as sticks. The MEK C-lobe is shown as a cyan surface and electrostatics surface (B). (C) The potential interaction of the B-Raf residue N661 with the indicated MEK residue shown as sticks. (D) The potential interaction of the B-Raf residue R671 with the indicated MEK residue shown as sticks. Hydrogen bonds and salt-bridges are shown with dashed lines and distances are provided. Figure generated using PDB ID: 4MNE ^*8*^.

Each mutant was expressed with a GST-tag in *E. coli* and fractions corresponding to dimer from size exclusion chromatography were isolated. We tested binding by BLI as described above. In agreement with our hypothesis, all mutants tested, except for R671A and the non-disruptive N661D mutant, had a higher K_D_ than the B-Raf^V600E,15mut^ construct (Table 1). Of note, the I666R and N661V mutations resulted in the greatest decrease (more than 10-fold) in binding affinity. This decrease was a result of both a slower association rate and an increased dissociation rate.

Given that B-Raf must bind to MEK before phosphorylating it, we next wanted to determine if the mutations in the B-Raf^V600E,15mut^ α-helix (residues 661-671) affect its kinase activity. To do this, we used the same *in vitro* phosphorylation assay as described above using GST-B-Raf constructs. Of all the mutants tested, only B-Raf^V600E,15mut, N661V^ and B-Raf^V600E,15mut,I666R^ showed significantly reduced levels of MEK phosphorylation when compared to B-Raf^V600E,15mut^, and the B-Raf^V600E,15mut,I666R^ mutant kinase was completely dead (Figure 6). The B-Raf^V600E,15mut,F667Y^ mutant showed a modest increase in activity under the conditions of the assay. As expected, B-Raf^V600E,16mut^ showed negligible activity, and MEK on its own was not able to autophosphorylate, so any phosphorylation observed was due to B-Raf. These data suggest that altering MEK binding by targeting the α-helix (residues 661-671) can alter MEK phosphorylation levels.

**Figure 6.**
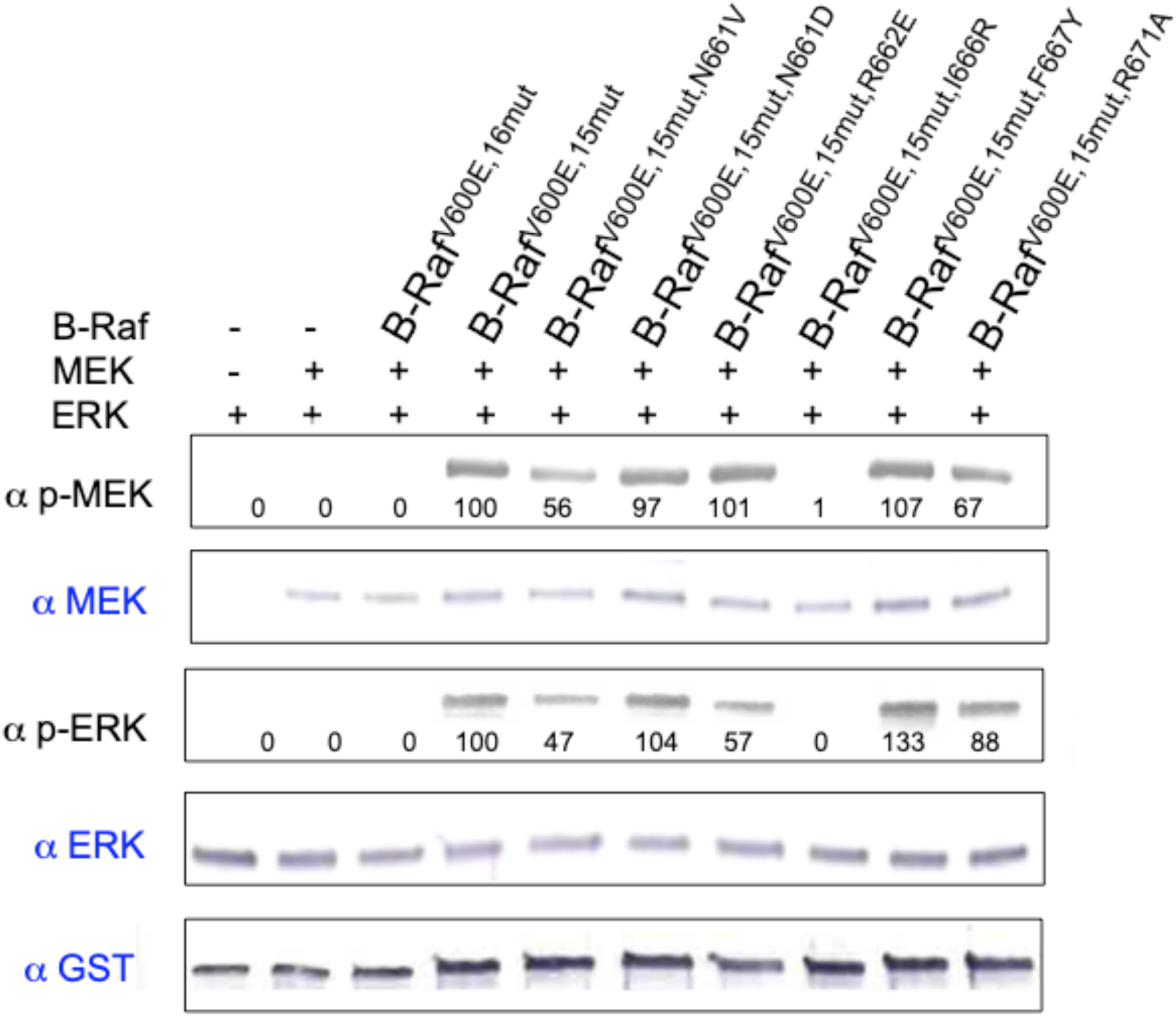
*In vitro* Phosphorylation of MEK and ERK by B-Raf ^V600E,15mut^ α-helix mutants. MEK and ERK were incubated with ATP with or without the indicated constructs of B-Raf for 30 minutes. Reaction mixtures were subjected to Western blot analysis at least twice with the indicated antibodies and a representative blot is shown. Input lanes are shown in blue. B-Raf and ERK were both GST-tagged. The intensities of the p-MEK and p-ERK bands were quantified by ImageJ and normalized to the corresponding band from the mixture with B-Raf ^V600E,15mut^.

### B-Raf α-Helix Point Mutations Affect ERK Phosphorylation

Activated MEK goes on to phosphorylate ERK at residues Thr202/Tyr204, so we wanted to determine the downstream effects of the B-Raf^V600E,15mut^ α-helix mutants. We repeated the same *in vitro* phosphorylation assay and probed for phosphorylation of ERK1 at residues Thr202/Tyr204. Consistent with the levels of MEK phosphorylation observed, ERK phosphorylation was drastically reduced in reaction mixtures that had B-Raf^V600E,15mut, N661V^ and B-Raf^V600E,15mut, I666R^ when compared to the reaction mixture in which B-Raf^V600E,15mut^ was used. This time, however, we also observed a reduction in ERK phosphorylation in reactions with B-Raf^V600E,15mut,R662E^. B-Raf^V600E,15mut, F667Y^ led to greater downstream ERK phosphorylation under the conditions of the assay. B-Raf^V600E, 16mut^ led to negligible ERK phosphorylation, as expected, as did MEK without prior phosphorylation by B-Raf^V600E,15mut^. ERK on its own was not able to autophosphorylate, so any phosphorylation observed was due to phosphorylated MEK (Figure 6).

To quantify the kinase activities of the V600E-containing B-Raf constructs, we used an ELISA assay as described above to measure p-ERK levels and to obtain K_M_(MEK) values, summarized in Table 2and Figure S2. When compared to B-Raf^V600E,15mut^, we observed that K_M_(MEK) for B-RafV600E,15mut,N661V, B-RafV600E,15mut,R662E and B-RafV600E,15mut,I666R are higher (indeterminate in the case of B-Raf^V600E,15mut,I666R^), the K_M_(MEK) for B-Raf^V600E,15mut,F667Y^ is lower, and the K_M_(MEK) for B-Raf^V600E,15mut,N661D^ and B-Raf^V600E,15mut,R671A^ are similar, consistent with the data from Western analysis. These data, which are consistent with K_D_ values we obtained from BLI, suggest that mutating the α-helix (residues 661-671) can alter ERK phosphorylation levels by altering B-Raf affinity to MEK and MEK phosphorylation.

### The B-Raf I666R Mutation in the Context of the Full Enzyme Reduces Activity in Cells

From the data above, the I666R mutation appeared to be most effective at reducing B-Raf activity. Thus, we next wanted to determine if introducing this mutation had an effect in cells in the context of a more biologically relevant B-Raf without the N-terminal truncation and solubilizing mutations of the B-Raf^V600E,15mut,I666R^ construct employed in the *in vitro* studies. To test this, we introduced the I666R mutation in the full length B-Raf (B-Raf^FL,V600E,I666R^), which includes the N-terminal regulatory domain, and the V600E mutation, and compared its activity to that of wild-type B-Raf^FL,WT^ and constitutively active B-Raf^FL,V600E^ constructs. HEK293T cells were transfected with B-Raf and MEK DNA and allowed to express for 24 hours before cell lysis and Western blot analysis of MEK and ERK phosphorylation levels. Consistent with our *in vitro* data, B-Raf^FL,V600E,I666R^ had lower activity than the constitutively active mutant B-Raf^FL,V600E^, but higher activity than B-Raf^FL,WT^ (Figure 7).

**Figure 7.**
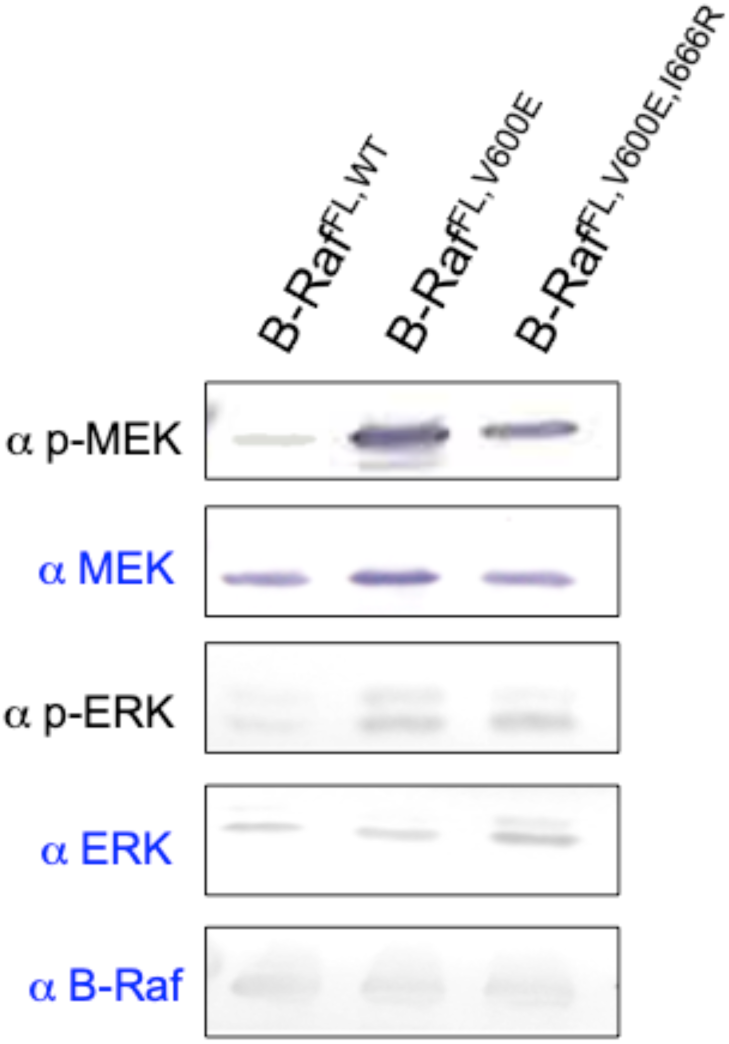
Phosphorylation of MEK and ERK by full length B-Raf in cells. Indicated full length B-Raf and MEK were transfected into HEK293T cells and lysed after 24 hours of expression. Cell lysates were subjected to Western blot analysis with the indicated antibodies. Control lanes are shown in blue. Experiment was performed at least three times and representative blots are shown.

## Discussion

A crystal structure of MEK1 in complex with the catalytic domain of wild-type B-Raf was determined by others to help understand B-Raf activity, important for MAPK signaling ^*8*^. Analysis of its structure revealed a residue that had been mutated in the solubilized construct of B-Raf for *E. coli* expression could make an important contact with MEK, thereby affecting its activity ^*31*^ (Figure 1B-1C). We showed that this mutation, when reverted to its wild-type residue (F667), restored MEK binding and phosphorylation activity in the *E. coli* construct (Figure 3A-3C, Figure 7). Our phosphorylation assay data revealed that B-Raf^V600E,15mut^ produced in *E. coli* had similar kinase activity as the wild-type B-Raf^V600E^ produced in Sf9 insect cells (Figure 3C). This is consistent with our finding that B-Raf^V600E,15mut^ had a similar K_D_ in binding MEK (∼32 nM) as B-Raf^V600E^ that was expressed in insect cells (K_D_∼56 nM). Also, we observed lower affinity and activity in our *in vitro* experiments for B-Raf^WT^. Structures have shown that the αC helix, which can form a salt bridge with a negatively charged moiety at residue 600 in the “active” conformation, is in a different orientation in the active and inactive conformations of B-Raf ^*8*^, which could affect MEK binding. Thus, our data with B-Raf^WT^ is consistent with data others obtained showing that B-Raf inhibitors that shift the αC helix toward an inactive conformation weaken MEK binding^*8*^. Still, a similar level of activity as B-Raf^V600E^ was observed for our *E. coli* B-Raf^V600E,15mut^ at saturating ATP concentrations, suggesting it is sufficient for studying the physiological functions of the constitutively active B-Raf^V600E^, particularly its interactions and phosphorylation of MEK and downstream effects on ERK. Our mutant, which can be expressed in *E. coli*, would also be useful for studies requiring large-scale protein preparations, such as those needed for drug screening efforts.

While current B-Raf inhibitors successfully prevent the progression of B-Raf^V600E^-related cancers for some time, inhibitor resistance and paradoxical activation can occur ^*25*^. Thus, we sought to explore a region of B-Raf that could be used as a drug target for disrupting B-Raf-MEK interactions. B-Raf residue F667, which we demonstrated is important for binding to and phosphorylation of MEK (Figure 4,Table 1), resides on an α-helix that makes extensive interactions with MEK (Figure 5A). We probed the importance of this α-helix by introducing mutations to alter binding to MEK. N661 appears to hydrogen bond with R234 of MEK and mutation to a hydrophobic residue (Val) would eliminate this interaction (Figure 5C). Mutation to Asp, however, could either improve binding by the introduction of a salt bridge, or maintain the same interaction as N661. R662 contacts an acidic patch of MEK, so its charge reversal to Glu would result in charge repulsion (Figure 5B). Similarly, I666 sits near a hydrophobic patch of MEK, so introducing a charged residue (Arg) would lead to an unfavorable interaction (Figure 5B). Finally, R671 appears to hydrogen bond with D315 of MEK, so its mutation to Ala would disrupt this interaction (Figure 5D). Our binding and kinetic data obtained by BLI and ELISA, respectively, are consistent with these hypotheses since the K_D_ and K_M_(MEK) values were reduced for all mutants predicted to disrupt binding (N661V, R662E, and I666R) except for R671A which had no effect. In the case of the N661D mutation, binding to MEK was also not affected, suggesting that the mutation to Asp maintains the hydrogen bond with R234 of MEK (Figure 5C). The F667Y mutation, however, was introduced to increase binding and activity, which we observed (Figure 4B,Figure 6,Table 1,Table 2). In addition to identifying a new target for drug discovery, the various interactions we highlight between the B-Raf α-helix (residues 661-671) and MEK might suggest useful properties (e.g. charges, hydrogen bond partners, and/or hydrophobic interactions) to take into account when designing inhibitors against this site.

Our *in vitro* and cell-based phosphorylation assays are consistent with our binding data (Figure 6). While some of the effects were not directly observed when probing for MEK phosphorylation, as in the case of the B-Raf^V600E,15mut, R662E^ mutant, it could be that the reaction time allowed even weaker MEK binders to complete phosphorylation. The effects of decreased activity of the B-Raf^V600E,15mut,R662E^ mutant were, however, apparent when probing for phosphorylated ERK. Our results suggest that the B-Raf-MEK α-helix (residues 661-671) interface is a promising target for drug discovery, as it is a different site from current B-Raf inhibitors. This was further illustrated in our cell-based assay using more biologically relevant B-Raf constructs. Of note, the fact that B-Raf^FL,V600E,I666R^ showed reduced activity when compared to B-Raf^FL,V600E^ suggests that targeting this site in cells could be effective against both B-Raf and B-Raf^V600E^-induced malignancies. B-Raf^FL,V600E,I666R^ did, however, have higher activity than B-Raf^FL,WT^, likely because heterodimers with C-Raf can still form and activate MEK ^*29*^. This result indicates that combination therapies with ATP-competitive inhibitors would be useful to further reduce B-Raf activity and prevent the development of resistance.

## Acknowledgements

D.F. acknowledges support from Swarthmore College Startup and Faculty Research Funds, R.M. acknowledges support from NIH grant CA226888, and L.A.A. acknowledges support from NIH grant CA226888-S1.

## Abbreviations

MAPK: mitogen-activated protein kinase
ATP: adenosine triphosphate
GTP: guanosine triphosphate
DTT: dithiothreitol
ELISA: enzyme-linked immunosorbent assay
FPLC: fast protein liquid chromatography
SEC: size exclusion chromatography

## Accession Codes

Proteins described in this study are in Uniprot, with the following accession codes: B-Raf (P15056), MEK (Q02750), ERK (P27361)

