## Supplemental Figures for "Identification and Characterization of a B-Raf Kinase Alpha Helix Critical for the Activity of MEK Kinase in MAPK Signaling"

*** Correspondence:**

Daniela Fera

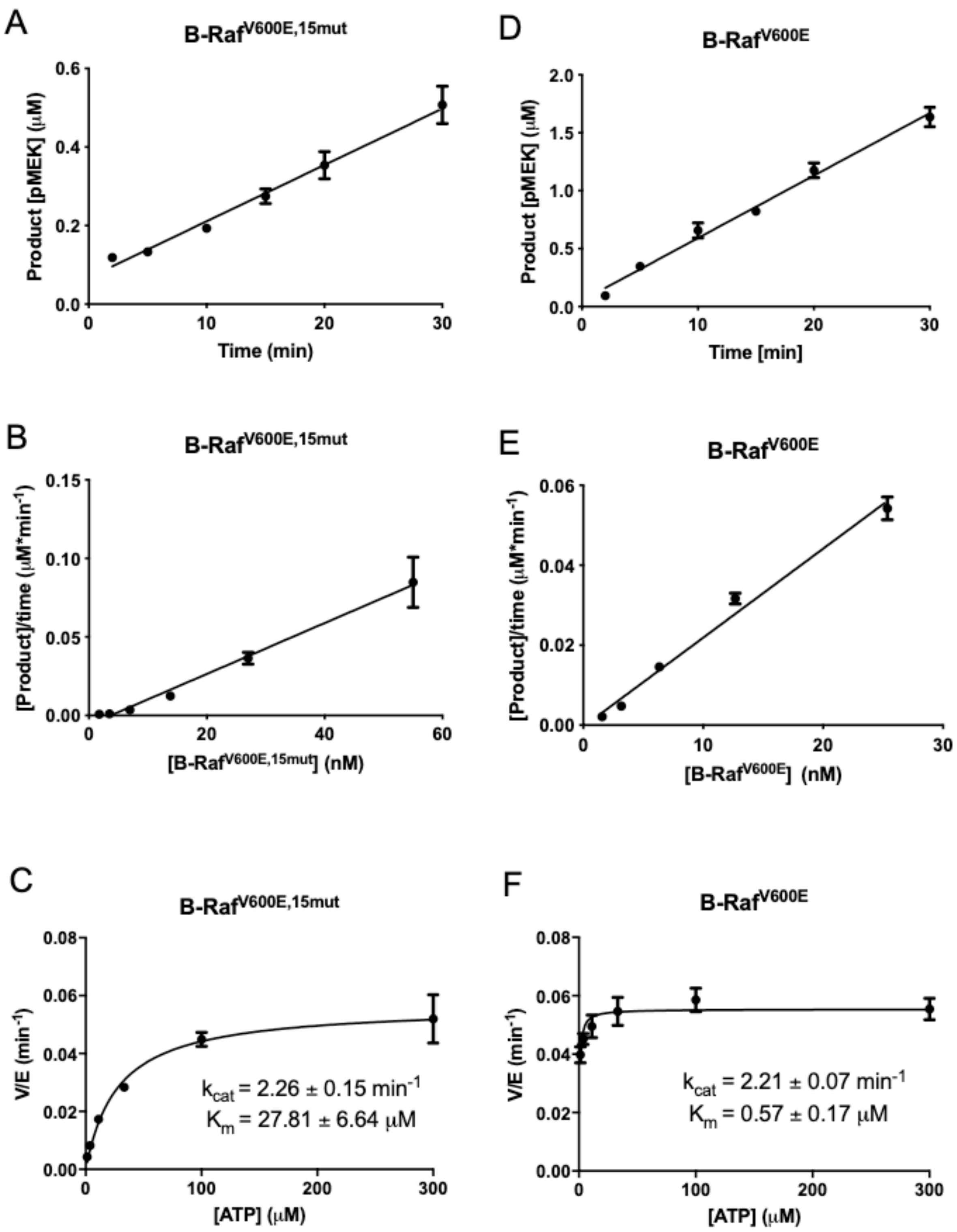


**Figure S1. Enzymatic characterization of B-Raf $^{V600E,15mut}$ and B-Raf$^{V600E}$ by ELISA.** The kinase activity of B-Raf $^{V600E,15mut}$ is linear versus (A) reaction time and (B) enzyme concentration. (C) Steady-state kinetic analysis of B-Raf $^{V600E,15mut}$. $K_m$ for ATP and $k_{cat}$ were obtained with varying concentrations of ATP at a fixed MEK substrate concentration (600 nM) and fixed B-Raf concentration (25 nM). The kinase activity of B-Raf$^{V600E}$ is linear versus (D) reaction time and (E) enzyme concentration. (F) Steady-state kinetic analysis of B-Raf$^{V600E}$. $K_m$ for ATP and $k_{cat}$ were obtained with varying concentrations of ATP at a fixed MEK substrate concentration (600 nM) and fixed B-Raf concentration (25 nM).

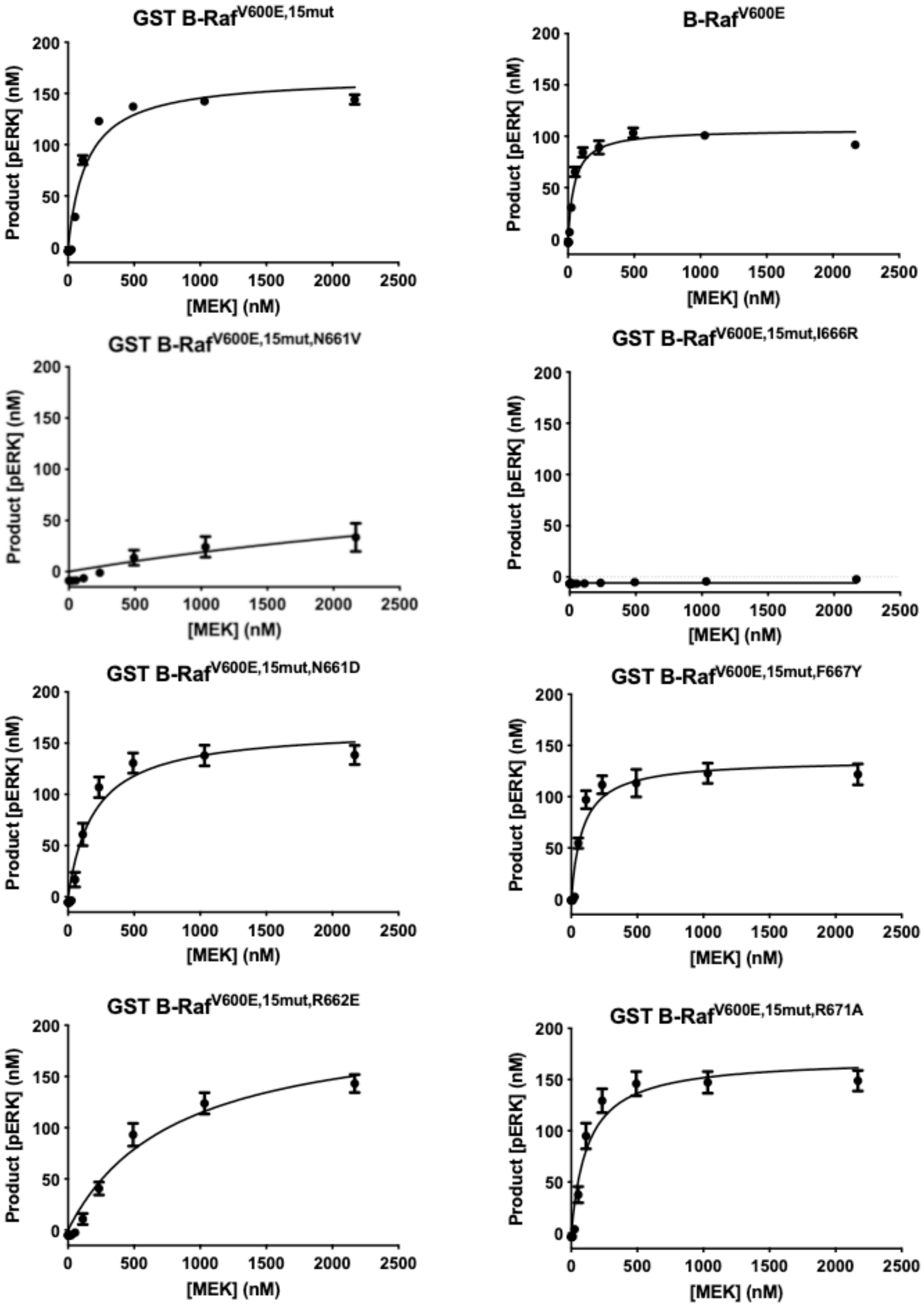
GST B-Raf$^{V600E,15mut}$
B-Raf$^{V600E}$
GST B-Raf$^{V600E,15mut,N661V}$
GST B-Raf$^{V600E,15mut,I666R}$
GST B-Raf$^{V600E,15mut,N661D}$
GST B-Raf$^{V600E,15mut,F667Y}$
GST B-Raf$^{V600E,15mut,R662E}$
GST B-Raf$^{V600E,15mut,R671A}$
Product [pERK] (nM)
[MEK] (nM)
0
50
100
150
200
0
500
1000
1500
2000
2500

**Figure S2. Enzymatic characterization of steady state parameters for MEK substrate of B-Raf$^{V600E,15mut}$ mutants and B-Raf$^{V600E}$ by ELISA.** $K_M$ (MEK) values were obtained for the indicated mutants with varying concentrations of MEK at a fixed ERK concentration (4.68 uM) and fixed B-Raf concentration (25 nM). Data for each mutant were obtained at least in duplicate.
